# *APOE4* alters the lipid droplet proteome and modulates droplet dynamics

**DOI:** 10.1101/2024.12.16.628710

**Authors:** Cassi M. Friday, Isaiah O. Stephens, Cathryn T. Smith, Sangderk Lee, Diksha Satish, Nicholas A. Devanney, Sarah Cohen, Josh M. Morganti, Scott M. Gordon, Lance A. Johnson

**Affiliations:** Department of Physiology, University of Kentucky; Sanders Brown Center on Aging, University of Kentucky; Department of Cell Biology & Physiology, University of North Carolina; Department of Neuroscience, University of Kentucky; Saha Cardiovascular Center, University of Kentucky

## Abstract

Excess lipid droplet (LD) accumulation is associated with several pathological states, including Alzheimer’s disease (AD). However, the mechanism(s) by which changes in LD composition and dynamics contribute to pathophysiology of these disorders remains unclear. Apolipoprotein E (ApoE) is a droplet associated protein with a common risk variant (E4) that confers the largest increase in genetic risk for late-onset AD. E4 is associated with both increased neuroinflammation and excess LD accumulation. In the current study, we sought to quantitatively profile the lipid and protein composition of LDs between the ‘neutral’ E3 and risk variant E4, to gain insight into potential LD-driven contributions to AD pathogenesis. Targeted replacement mice expressing human E3 or E4 were injected with saline or lipopolysaccharide (LPS), and after 24 hours, hepatic lipid droplets were isolated for proteomic and lipidomic analyses. Lipidomics revealed a shift in the distribution of glycerophospholipids in E4 LDs with a concomitant increase in phosphatidylcholine species, and overall, the baseline profile of E4 LDs resembled that of the LPS-treated groups. Quantitative proteomics showed that LDs from E4 mice are enriched for proteins involved in protein/vesicle transport but have decreased levels of proteins involved in fatty acid β-oxidation. Interestingly, proteins associated with LDs showed substantial overlap with previously published lists of AD postmortem tissue and microglia ‘omics studies, suggesting a potential role for LDs in modulating AD risk or progression. Given this, we exposed primary microglia from the same E3 or E4 mice to exogenous lipid, inflammatory stimulation, necroptotic N2A cells (nN2A), or a combination of treatments to evaluate LD formation and its impact on the cells’ immune state. Microglia from E4 mice accumulated more LDs in every condition tested – at baseline and following addition of fatty acids, LPS stimulation, or nN2As. E4 microglia also secreted significantly more cytokines (TNF, IL-1β, IL-10) than E3 microglia in the control, oleic acid, and nN2A treatment conditions, yet showed a blunted response to LPS. In sum, these results suggest that E4 microglia accumulate more LDs compared to E3 microglia and that E4 is associated with a basal LD composition that resembles a pro-inflammatory cell. Together with the high overlap of the LD proteome with established AD-associated datasets, these data further support the idea that alterations in LD dynamics, particularly within microglia, may contribute to the increased risk for AD associated with *APOE4*.

## INTRODUCTION

Lipid droplets (LDs) are ubiquitous storage organelles which are conserved across many species and expressed in multiple cell types.^1-3^ Although historically presumed to be innocuous and inert lipid storage depots, many studies now highlight the dynamic nature of LDs in terms of their formation, turnover, and composition across a number of homeostatic and pathological conditions, including neurodegenerative disease.^4^ Both the lipid content and outer protein profile of an LD can vary greatly, with wide-ranging consequences for cellular function. For example, LDs liberate fatty acids as lipid mediators of inflammation^5^ or for mitochondrial metabolism.^6^ Differences in either the inner neutral lipid content or the phospholipid-rich shell can influence the assembly and function of proteins on the LD monolayer.^7^

A set of ubiquitous LD proteins known as perilipins (PLINs) provide clues about LD function based on their expression within the cell. For example, PLIN5 is associated with oxidative phosphorylation and colocation of LDs to the mitochondria,^3,8-11^ whereas PLIN2 expression is associated with immunologic and inflammatory function within the cell.^12-13^ With this information in mind, it is reasonable to believe the LD composition and protein makeup is as dynamic as their now recognized multitude of functions and should be considered when studying LD accumulation in disease. It is well documented that several diseases are associated with excessive LDs, including but not limited to obesity, atherosclerosis, cancer, type 2 diabetes, and Alzheimer’s disease (AD);^4,14^ and evidence suggests these organelles may be active participants in disease mitigation, progression, or both.^15^

Genome-wide association studies (GWAS) identify many late-onset Alzheimer’s disease (LOAD) risk genes involved in the immune response and microglia function,^16^ particularly genes involved in microglial phagocytosis and debris processing.^17^ The strongest of the LOAD risk genes is the ε4/ε4 (E4) allele of Apolipoprotein E (*APOE*), a gene also strongly implicated in microglial function.^18-20^ *APOE* is a disease-associated-microglia (DAM) gene and diseased or stressed microglia upregulate production of ApoE.^18^ Intriguingly, ApoE is also droplet-associated protein found on the LD surface,^21-22^ and E4 expressing astrocytes^23-25^ and E4 microglia^26-28^ both accumulate more lipid droplets than neutral AD-risk ε3/ε3 (E3) expressing cells. Additionally, carriage of the E4 allele is associated with increased neuroinflammation and metabolic disturbances within neurons and glial cells,^18^ two functions potentially altered by LD accumulation and composition.

It is of particular interest that glial cells accumulate LDs in response to neuronal stress, age, or neurodegeneration.^4,30-31^ LD accumulation has been described as both a beneficial compensatory response and a pathological feature within the brain. For example, there is compelling evidence of a synergistic relationship between neurons and astrocytes whereby fatty acids are shuttled from neurons to sequester in astrocytic LDs to protect neurons against fatty acid toxicity.^30^ However, a more pathologic accumulation of LDs associated with age in microglia causes abnormal cellular function.^30^ Lipid accumulation with age is highly relevant in Alzheimer’s disease (AD), where even the first description of AD by Alois Alzheimer noted glia with adipose inclusions.^32^

While there are numerous reports of both peripheral and CNS cell types accumulating more LDs in the presence of E4, the effects of *APOE* on the composition of LDs themselves remains unexplored.^24-29,33-34^ Here, we sought to develop a deeper knowledge of LD characteristics between *APOE3 and APOE4* expressing cells in the liver and brain. E3 and E4 expressing LDs were analyzed for their relative accumulation at baseline and in response to an inflammatory challenge, their physical characteristics (including size, lipid, and protein composition), and their potential effects on microglia function.

## RESULTS

### LD-enriched fractions show greater droplet accumulation in E4 and inflamed conditions

Extracting sufficient LD from microglia – either *in vitro* or *in vivo* – is prohibitive due to their relatively small LD content compared to the large quantities of cellular material necessary for successful LD isolation and quantitative ‘omics. Therefore, we first studied the more abundant and easily accessible LDs from the liver of human E3 and E4 expressing mice to quantify ApoE isoform-specific changes in LD composition (Fig. 1a). Mice were injected with saline or LPS (5mg/kg) and liver tissue was harvested after 24 hours. LDs were enriched via density gradient centrifugation where the buoyant, lipid-rich fractions at the top of the centrifuge tube were collected for western blot verification of lipid droplet surface protein PLIN2 (Fig. 1b). Consistent with previous studies,^33^ Oil Red O staining showed higher lipid content in E4 liver tissue compared to E3 at baseline (Fig. 1c-e). Both E3 and E4 liver tissue increased lipid content after LPS stimulation, although this response was muted in the E4 LPS compared to E3 LPS livers (Fig. 1c-d). Together, these data validate an *APOE*-specific, LD-enriched source material for downstream analysis of the proteome and lipidome.

**Figure 1.**
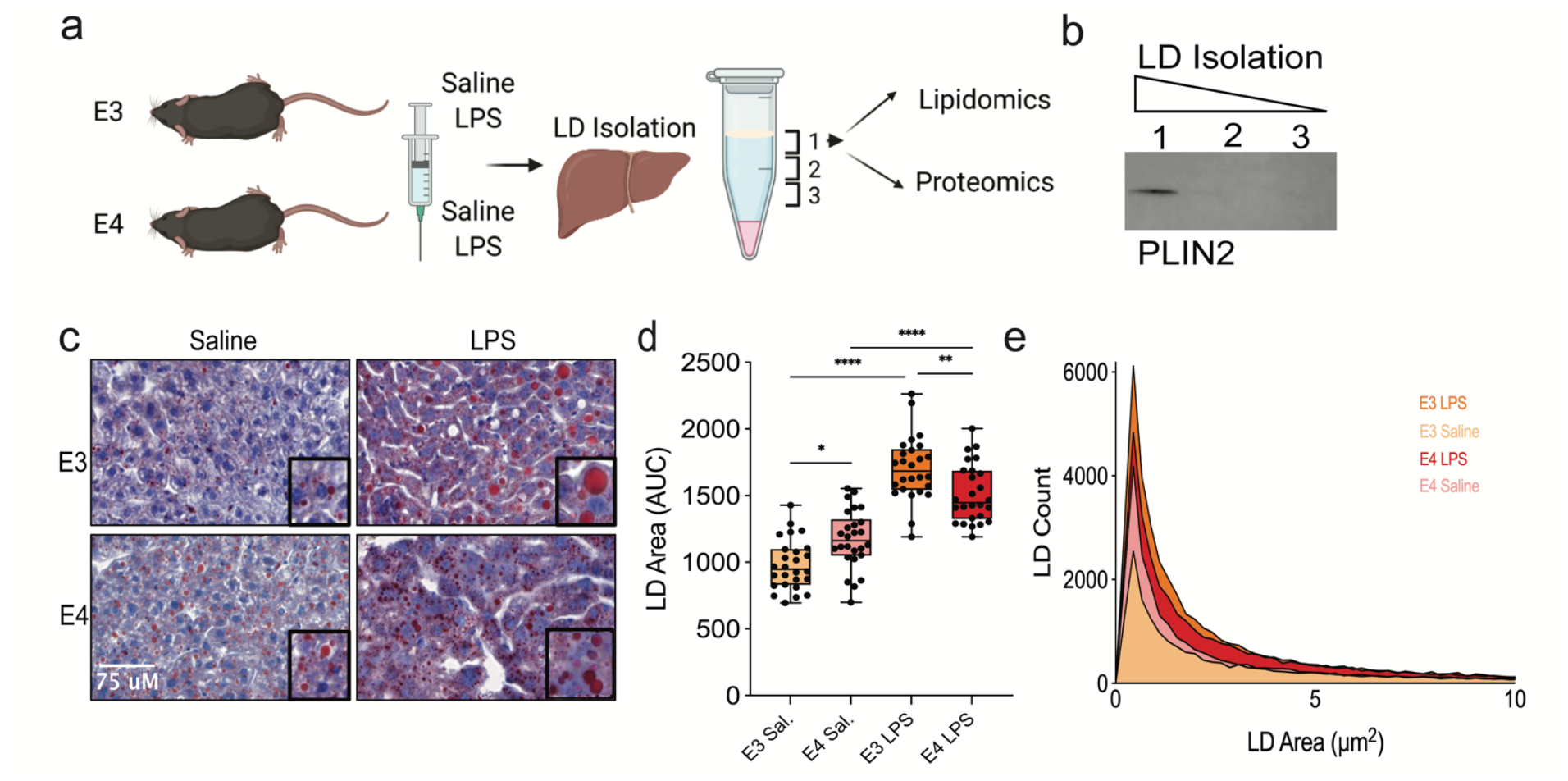
Lipid droplet isolation from mouse liver tissue successfully targeted the droplet fraction and inflamed liver tissue showed increased lipid via ORO stain. a) Targeted replacement mice expressing human ApoE3 or ApoE4 (n = 5/group) were either injected with saline or 5mg/kg of LPS. After 24 hours, liver tissue was homogenized for lipid droplet enrichment. b) Western blot of the top sequential liquid volumes after density gradient centrifugation showed LD marker, PLIN2 in only the top fraction. This fraction was analyzed for proteomics and lipidomics. c) ORO staining of liver sections from the same mice to quantify lipid amount (n=3, 9 images analyzed per mouse). d) Quantification of ORO staining represented as total lipid area. e) Histogram showing size distribution of LDs analyzed from ORO staining.

### E4- and LPS-stimulated LDs are enriched in phosphatidylcholine

As anticipated, the bulk of lipid content in LD enriched fractions was composed of glycerolipid, with ∼90% of the lipidome consisting of triacylglycerols (TGs), a common neutral lipid component of the LD inner core (Fig. 2a, Supplemental Table).^36^ The other lipid species detected were largely neutral diacylglycerols (DGs) and glycerophospholipid species that make up the outer LD monolayer (Fig. 2a, large circles). Phosphatidylcholine (PC) was notably enriched in E4 compared to E3 LDs at baseline, as well as in both LPS treated groups (Fig. 2a). Similarly, volcano plots of differentially abundant metabolites show that while E3 droplets increase their PC content when stimulated with LPS, a similar response is not observed in E4 droplets (Fig. 2b). These results suggest that unlike E3 LDs, which substantially remodel their lipidome in response to an inflammatory challenge, E4 LDs show a blunted response and display a baseline lipid profile that already mirrors the post-inflammatory state.

**Figure 2.**
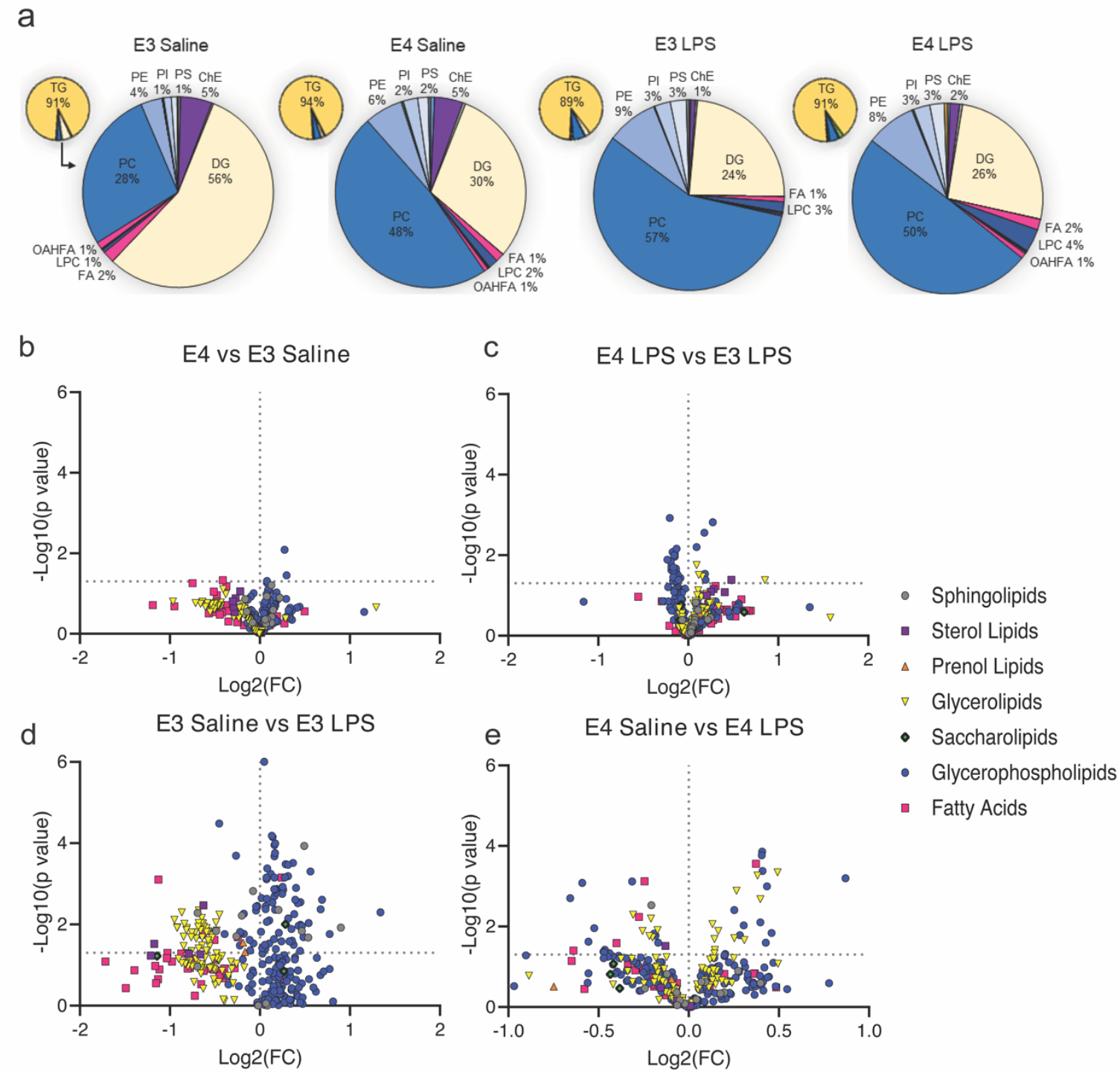
The lipidome of E4 droplets at baseline is similar to LPS-treated droplet lipidomes. a) The small pie charts indicate the majority abundance of triacylglycerol in the droplet lipidomes. The larger pie chart expands the remaining lipid species to observe lipid distributions between genotypes and treatments. b) Volcano plot of differentially abundant lipids by class between baseline conditions (higher abundance in E4 saline on the right). c) Volcano plot of differentially abundant lipids by class between E3 LPS and E4 LPS treatments. d-e) Volcano plots of differentially abundant lipids by class within the same genotype from baseline (saline injection) to LPS treatment

### E4 droplets are enriched with TG sequestration proteins, depleted in fatty acid β-oxidation proteins

Quantitative proteomic analyses of the LD-containing fractions revealed at total of 4,638 LD-enriched proteins (Supplementary Table). We cross referenced these with other known LD proteomes to identify proteins that are consistently and/or exclusively expressed on the LD surface (Fig. 3a, Supplementary Table), which resulted in a total of 2,338 ‘LD resident’ proteins which overlapped with at least one other dataset.^12,22,37-40^ Gene ontology analyses showed that under basal conditions, E4 LDs were enriched for proteins involved in LD organization, membrane trafficking, and TG sequestration, while they were relatively depleted in proteins related to fatty acid β-oxidation (Fig. 3b). Following LPS treatment, a comparison of the proteome between E4 and E3 LDs reflected differences in the type of immune proteins localized to the LD surface (Fig. 3c). Finally, a within-genotype comparison of the LPS response highlighted changes in LD proteins related to reorganization and transport, as well as several metabolic pathways (Fig. 3d-e).

**Figure 3.**
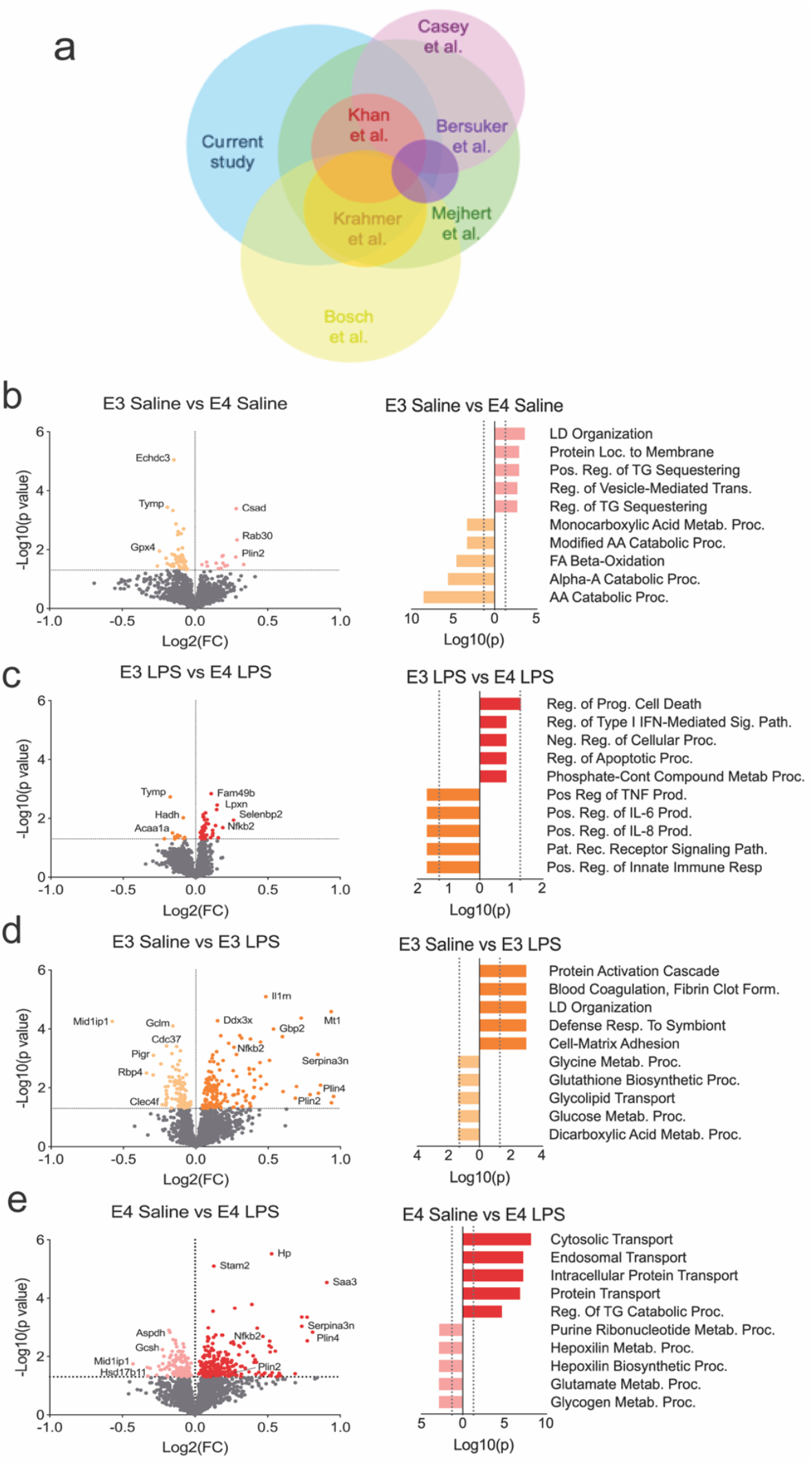
Differentially expressed proteins and gene ontology reveals enriched metabolic and immune pathways on E3 and E4 LDs. a) Venn diagrams showing overlap between the APOE LD proteome (“Current study”) and previously published studies. b) Gene ontology from differentially expressed proteins on E3 droplets compared to E4 at baseline highlight LDs and fatty acid beta-oxidation. Volcano plot of differentially expressed proteins between E3 and E4 droplets at baseline. c) Gene ontology from differentially expressed proteins between E3 and E4 droplets after LPS stimulation highlights apoptotic pathways and the innate immune response. Volcano plot of differentially expressed proteins between E3 and E4 droplets after LPS stimulation. d) Gene ontology from differentially expressed proteins between E3 saline and E3 LPS LDs. Volcano plot of differentially expressed proteins between E3 saline and E3 LPS LDs. e) Gene ontology from differentially expressed proteins between E4 saline and E4 LPS LDs. Volcano plot of differentially expressed proteins between E4 saline and E4 LPS LDs. AA, amino acid; FA, fatty acid; Loc, location; Neg, negative; Pat, pattern; Pos, positive; Proc, process; Rec, receptor; Reg, regulation; TG, Triglyceride.

To identify potential protein networks, we next performed a weighted gene co-expression network analysis (WGCNA) of the LD proteome (Fig. 4a-b). Several modules were significantly up- or down-regulated by *APOE* genotype or LPS treatment, including the green-yellow module, which was down-regulated in E4 LDs and associated with fatty acid β-oxidation and amino acid metabolism (Fig. 4c). The brown and purple modules, which were upregulated by LPS in both genotypes, were enriched in pathways involved in host response and glycogen metabolism or protein and vesicle transport, respectively (Fig. 4d, Supplemental Figure 1). In sum, these results imply that i) the E3 vs E4 response to an inflammatory challenge features varying LD protein dynamics, and ii) the E4 LD is enriched for proteins that facilitate lipid storage and inhibit utilization.

**Figure 4.**
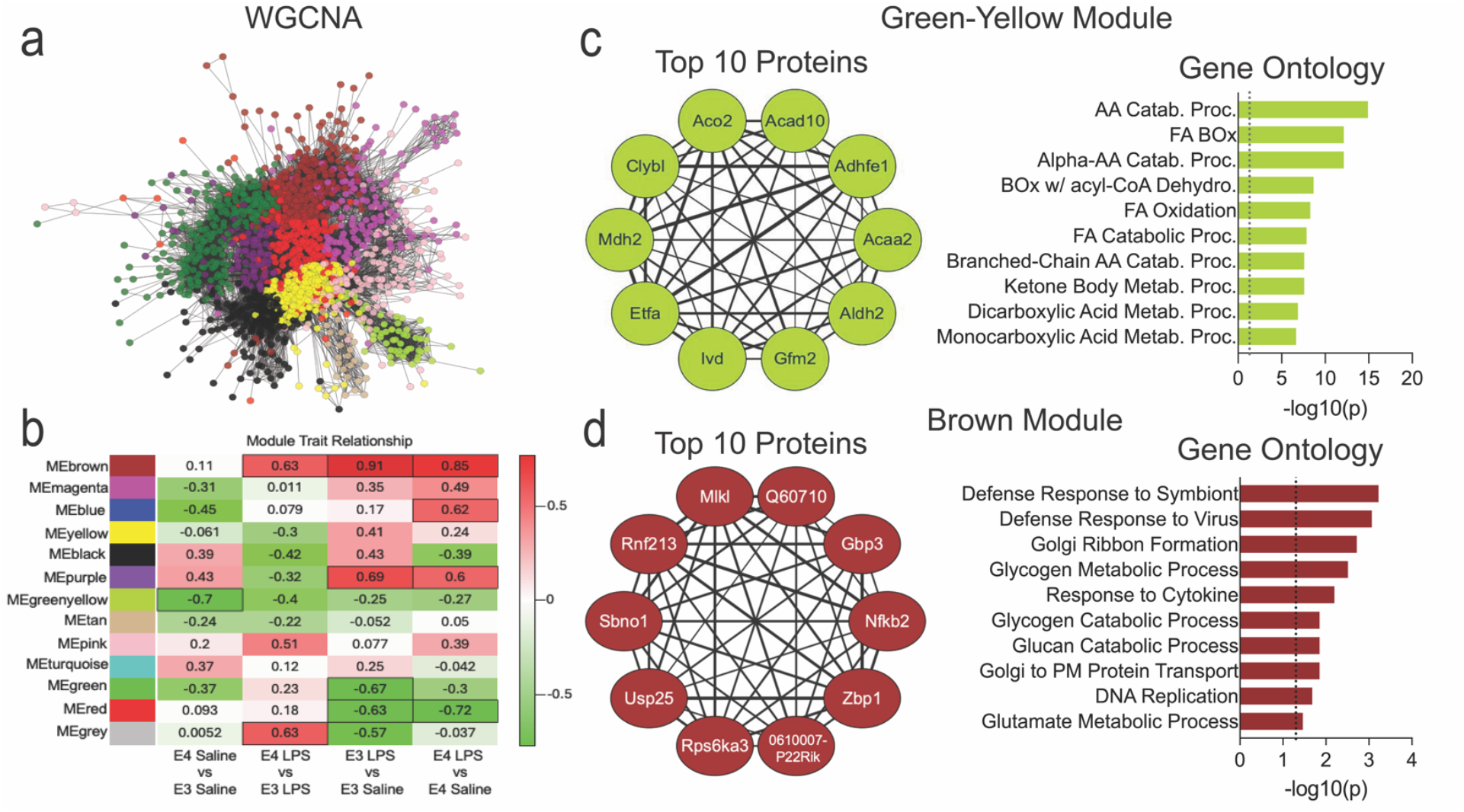
WGCNA analysis reveals significant difference in fatty acid metabolism and vesicle transport pathways between E3 and E4 Droplets. a) Whole network representation of our weighted gene correlation network analysis (WGCNA). b) Heatmap displaying all 13 WGCNA modules for each comparison. Boxes which are outlined represent significant difference with a p-value <0.05. c) The green-yellow module was one of the most significantly different between genotypes with E4 having lower enrichment for pathways involved in lipid metabolism. The 10 ten proteins from this module are displayed in the circle plot along with gene ontology form the entire module. d) The brown module was enriched in pathways involved in defense response and glycogen metabolism which had higher enrichment in E4 LPS. The top 10 genes from this module are displayed in the circle plot along with the gene ontology form the entire module. AA, amino acid; BOx, beta-oxidation; Catab., catabolism; FA, fatty acid; Proc., process.

### Proteins previously tied to *APOE4* and AD are largely LD-enriched

The LD proteome analyses above revealed several proteins that we recognized from the AD literature. To systematically assess this, we mined several proteomic and transcriptomic datasets for comparison, including a unique protein signature referred to as “incipient AD” (iAD).^41^ The iAD signature is a set of proteins highly expressed in young E4 carrier brains but reduced in AD brains.^41^ Intriguingly, 15 out of 25 (60%) of these AD-predictive proteins were highlighted in our LD proteome (Fig. 5a-b, Supplementary Figure 2). Another proteomic analyses of human AD brain identified a network module related to microglial metabolism (termed “M4”).^42^ In that study, Johnson et al characterized the top 30 most differentially expressed microglial transcripts from an AD mouse model that corresponded to M4. Notably, this list showed near complete overlap (27 of 30 proteins) with our LD proteome (Fig. 5c-d, Supplementary Figure 3). Similar results can be seen from a comparison with Dai et al where ∼35% of proteins upregulated in E4/4 AD compared to E3/3 AD brains^43^ overlapped with proteins enriched on E4 LDs. In contrast, ∼29% of proteins upregulated in E3/3 AD compared to E4/4 AD brains overlapped with proteins enriched on E3 LDs and were associated with pathways of fatty acid β-oxidation (Supplemental Table). We next overlaid our list of LD-enriched proteins with differentially expressed genes in microglia from E3- or E4-expressing mice treated with either saline or LPS.^19^ In both the saline and LPS treated conditions, the overlap of LD proteins and microglial genes differentially regulated by E4 pointed to an upregulation of pathways of protein synthesis and transport along with a concomitant downregulation of fatty acid metabolism (Fig. 5e-f). Together, these data show the protein composition of liver-derived LDs – and the effect of E4 in modulating enrichment of these proteins – substantially overlaps with the established E4/AD proteome, suggesting that many of the same proteins that define AD pathogenesis are found on or closely associated with LDs.

**Figure 5.**
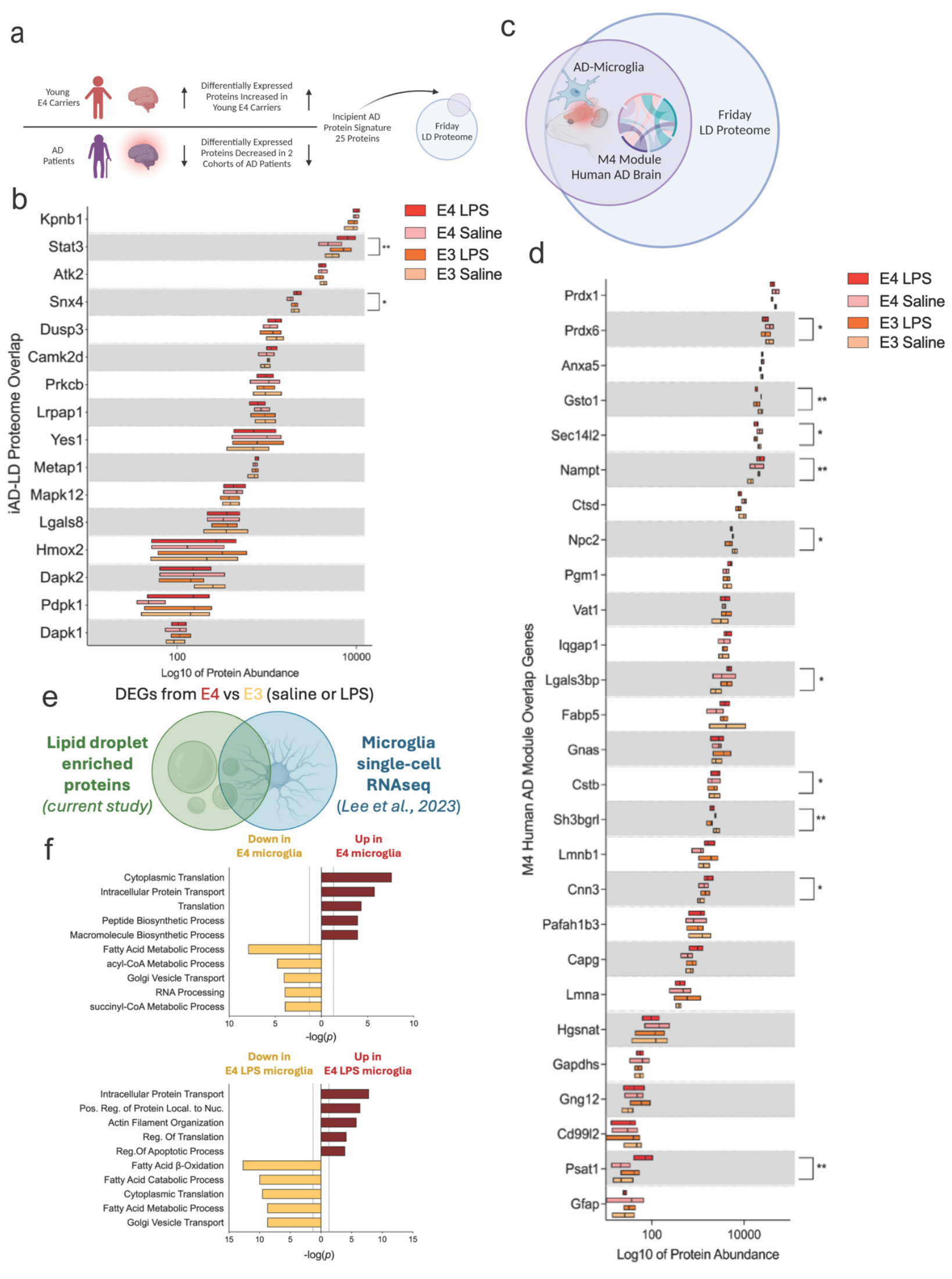
Human AD proteomes have significant overlap with LD proteins. a-b) A previous study by Roberts et al. identified a unique signature where proteins are increased in young E4-carriers compared to non-carriers but decreased in AD brains compared to cognitively normal brains. This “incipient AD” (iAD) signature showed 60% overlap with the APOE LD proteome from the current study. b) Box and whisker plot of overlapping iAD proteins. *p<0.05, **p<0.005, two-way ANOVA analyses. c-d) A large proteomic study by Johnson et al. revealed a glial metabolism module (“M4”) which was differentially expressed in AD brains as well as in microglia from a mouse model of AD. This AD-glial-metabolism signature has 90% overlap with our LD proteome. d) Box and whisker plots of the 27 overlapping AD-glial-metabolism proteins. *p<0.05, **p<0.005, two-way ANOVA analyses. e-f) LD-enriched proteins were compared with differentially expressed genes from microglia harvested from E3- or E4-expressing mice treated with either saline or LPS (Lee et al. 2023). f) Gene ontology of LD proteins which were also differentially expressed in E4 vs E3 saline (top) or LPS treated (bottom) microglia. Microglial LD genes differentially regulated by E4 were tied to an upregulation of pathways of protein synthesis and transport along with a concomitant downregulation of fatty acid metabolism.

### E4 microglia accumulate more LDs and secrete more cytokines following a variety of stimuli

Given the ostensible AD-microglia signature in the LD proteome, we examined LDs in E3 and E4 primary microglia. To test differences in LD accumulation between E3 and E4 microglia, we compared untreated microglia to those supplemented with exogenous lipid (oleic acid, OA), a pro-inflammatory stimulus (LPS), necroptotic N2A cells (nN2A), or a combination of these treatments. In each condition, E4 microglia showed an increase in LD accumulation compared to E3 (Fig. 6a-b). To determine the functional consequences of LD inhibition, we also treated microglia with either the DGAT1 inhibitor A922500 or Avisimibe, an ACAT1 inhibitor (Fig. 6c).^44-45^ As expected, the DGAT1 inhibitor prevented LD accumulation in all conditions with the exception of nN2A treated cells, where LDs can still form through the ACAT pathway (esterification of cholesterol) from neuronal debris (Fig. 6d). Similarly, the ACAT1 inhibitor prevented LDs in all conditions except those treated with OA, where synthesis of LDs can occur through the DGAT pathway (acetylation of DG to TG) (Fig. 6e).

**Figure 6.**
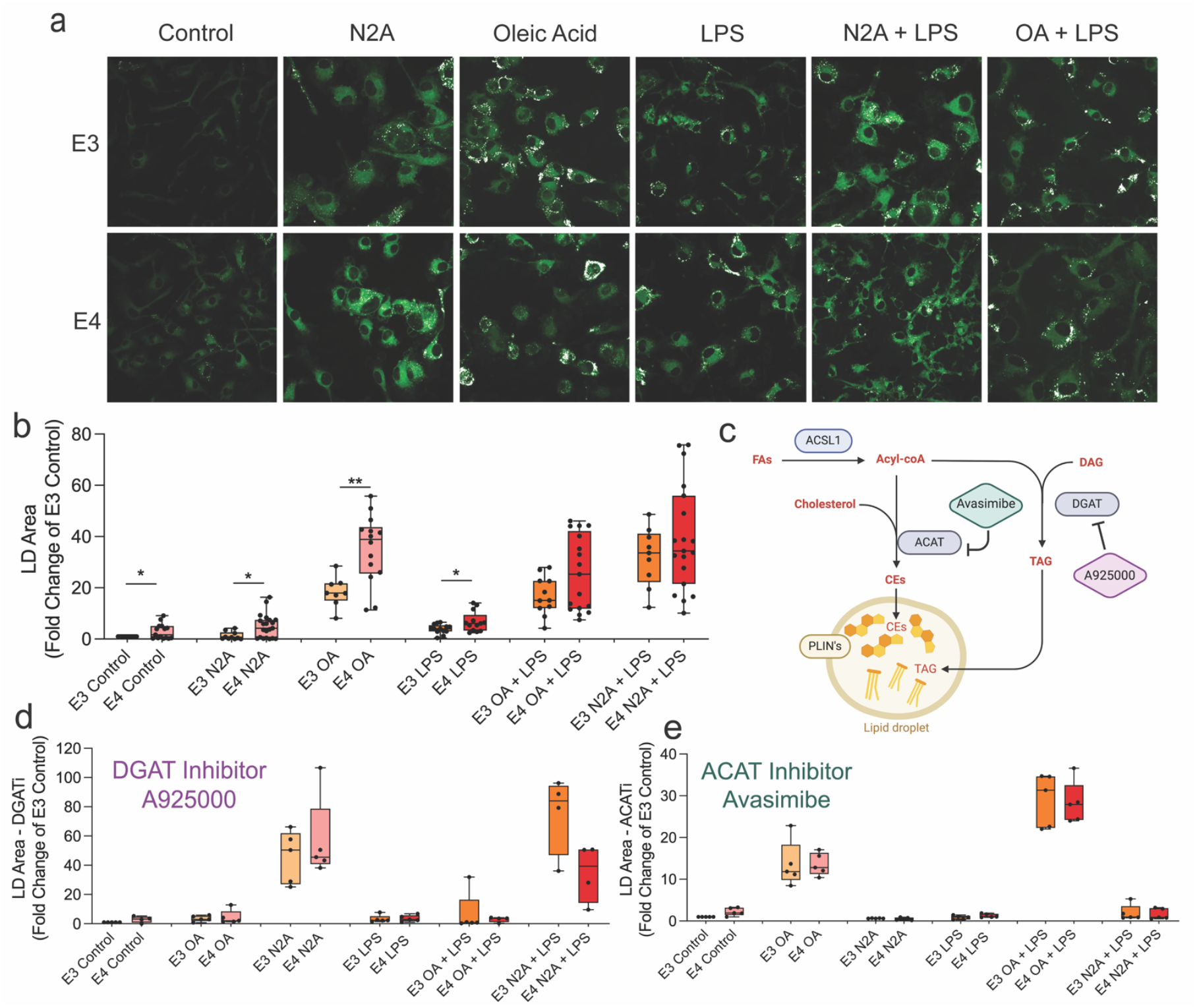
E4-expressing microglia accumulate more LDs than E3 microglia across multiple stimuli; inhibitors attenuate LD formation equally in E3 and E4 microglia. A-B) Primary microglia from E3 and E4 mice were treated with exogenous fatty acid (oleic acid (OA)), an inflammatory stimulus (LPS), necrotic neurons (nN2A), or a combination and LD formation was assessed 24 hours later. E4 microglia accumulated significantly more LD than E3 across control, OA, N2A, and LPS-treated conditions.. \**p*<0.05, \*\**p*<0.01, t-test. C) Schematic showing the inhibition of cholesterol ester (CE) formation via ACAT or inhibition of triacylglycerol (TAG) via DGAT. D) The DGAT1 inhibitor A925000 reduced droplet formation across all conditions with the exception of nN2A treatment and did so similarly in E3 and E4 microglia. E) The ACAT1 inhibitor Avasimibe prevented LD formation in both E3 and E4 microglia treated with either LPS or nN2As, but not as robustly in OA treated cells.

Lipids within LDs can be liberated to become mediators of inflammation, and in aging and AD, lipid droplets accumulate in microglia and cytokine production increases.^6,28,31^ Measurement of cytokine release into the media showed that E4 microglia secreted more TNF, IL-1β and IL-10 than E3 at baseline, as well as when exposed to OA and nN2A (Fig. 7a-d). As expected, the addition of LPS resulted in a dramatic increase in the release of these cytokines in control, OA, and nN2A treated cells; however, this effect was blunted in E4 microglia for TNF and IL-10 (Fig. 7e-g). Notably, while blocking ACAT1 lowered IFN-γ secretion, particularly in E4 microglia, preventing LD formation by inhibiting either ACAT1, DGAT or both did not decrease cytokine output in most conditions (Fig. 7a, Supplemental Table & Figures 4-5). Together, these data suggest that E4-expressing microglia accumulate more LDs and secrete higher levels of cytokines, findings in line with other recent studies.^26-28,46^ However, our results also suggest that – regardless of *APOE* genotype – inhibition of LD formation is largely insufficient to reduce cytokine production.

**Figure 7.**
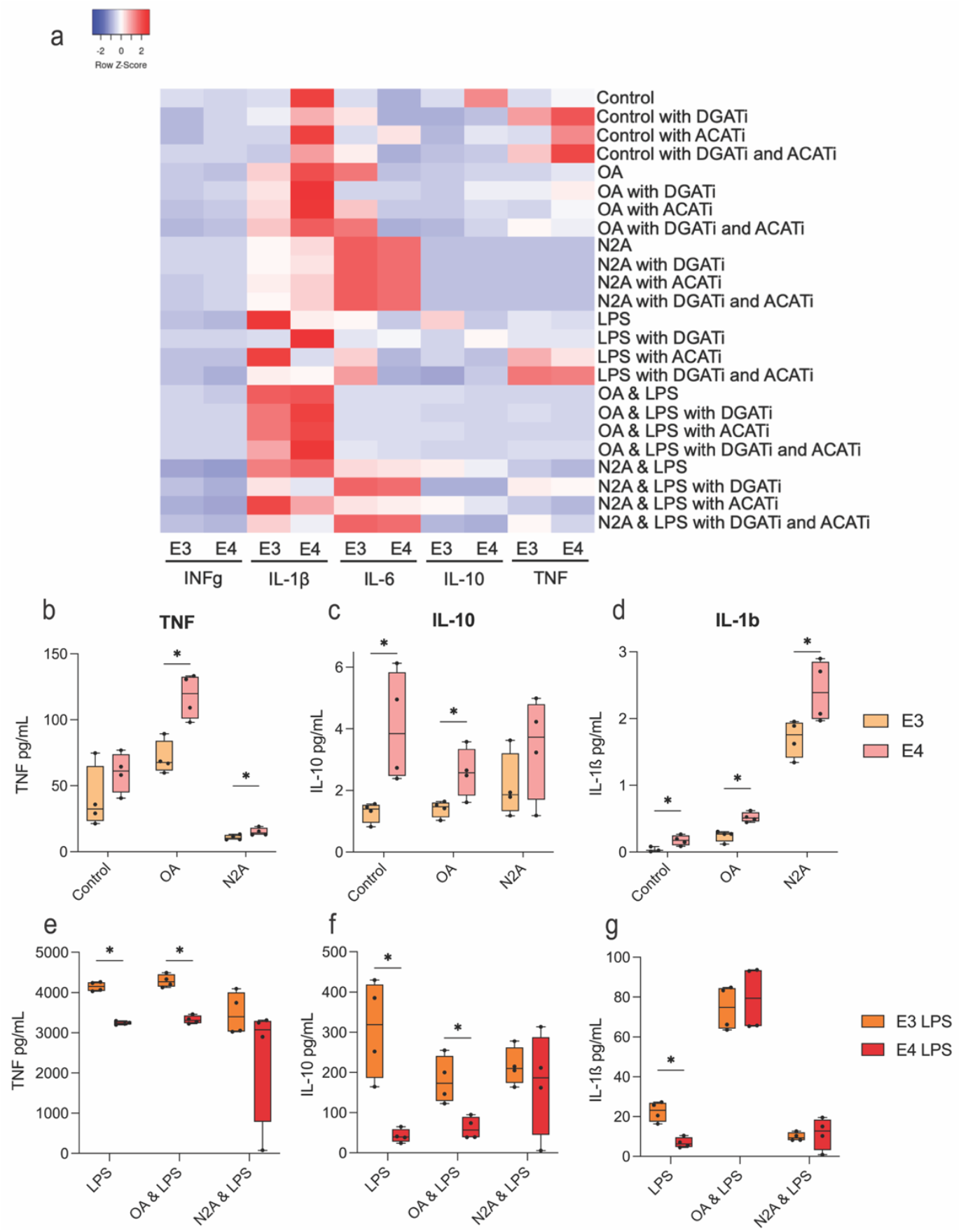
E4 microglia secrete more cytokines at ‘baseline’, but do not respond as robustly to LPS; inhibition of LD formation does not blunt cytokine release. A) Heat map showing cytokine levels in the media of E3 and E4 microglia following various stimuli (OA, LPS, or N2A) with or without inhibition of LD formation. B-D) Concentrations of TNF, IL-10, and IL-1ß measured in the media of E3 or E4 microglia after loading with OA or nN2A (i.e. ‘baseline’). E-G) Concentrations of TNF, IL-10, and IL-1ß measured in the media of E3 or E4 microglia after loading with OA or nN2A plus stimulation with LPS.

## DISCUSSION

In the current study, we provide the first *APOE* genotype-specific characterization of the lipid droplet-ome. We show that LDs from humanized E4 mice have a distinct lipid and protein profile compared to E3 droplets, both at baseline and following an inflammatory challenge. Building upon a growing literature, we also show that E4 expressing microglia accumulate more LDs and secrete more cytokines following various stimuli. Finally, by comparing our APOE-LD proteome with previous ‘omics analyses of postmortem AD/E4 brain and AD/E4 microglia, our data suggest that many of these previously implicated proteins are LD-associated.

Rather than inertly sequestering lipid, it is now appreciated that LDs play a much more active role within the cell, working to integrate metabolism and the immune response by hosting a number of different proteins on their membranes. One of these proteins recently shown to traffic to the LD surface is ApoE,^23^ an apolipoprotein whose common variants impart dramatically different risk for AD.^47-48^ Our findings of more LDs in E4-expressing microglia are in agreement with a growing number of studies showing a disproportionate buildup in E4 compared to E3 livers^33^, fibroblasts^49^, astrocytes^24-25,29,50^, and microglia^26-28^. Additionally, a number of studies have examined the effects of *APOE* genotype on the lipid profile of glia in vitro^23,25,29^. However, these previous studies did not address a possible effect of *APOE* on the composition lipid droplets themselves. In our view, the connection between ApoE itself as a droplet resident protein, ApoE’s role in lipid trafficking and metabolism, and E4’s role as the largest genetic risk factor for late-onset AD implored further investigation into LD dynamics between *APOE* genotypes.

Lipidomic analysis of our isolated LD fractions indicated that the droplets were primarily composed of TGs, a finding expected of droplets in the liver. However, other lipids within the droplet core and membrane differed markedly by genotype and treatment condition. At baseline, E4-LDs showed an enrichment in PC that mirrored the lipid profile of LPS-treated droplets. This finding could be indicative of a smaller droplet size and increased surface-to-volume ratio that would require increased phospholipids in the membrane. Specifically, PC incorporation in the LD monolayer has been shown to act as a surfactant to prevent LD coalescence,^51^ and an increased surface-to-volume ratio of smaller droplets may provide clues to their fate within the cell; i.e. larger droplets are more prone to lipolysis for fatty acid liberation to contribute to ß-oxidation, while smaller droplets are more prone to lipophagy.^52^ In line with this, our lab previously described smaller LDs in E4 astrocytes along with an E4-associated shift away from FA oxidation and toward aerobic glycolysis.^24,53^ This line of thinking is bolstered by the proteomics gene ontology and WGCNA results which showed a depletion of FA β-oxidation proteins on the E4 LDs relative to E3.

Regarding the LD proteome, an exciting finding from our profiling of LD-enriched liver fractions was a surprisingly high overlap with multiple AD/APOE4/microglia proteomic and transcriptomic datasets. For example, we found significant overlap of our LD-enriched protein list with an “incipient AD” proteomic signature of young E4-carriers published by Roberts et al.^41^ In this study, tissue from two human AD studies were analyzed for proteomic differences between cognitively normal and AD brains. Differentially expressed proteins from both studies were compared to significantly changed proteins in the brains of young E4-carriers and non-E4-carriers. This revealed a set of 25 proteins that were intriguingly upregulated in young E4 carriers yet downregulated in AD. Interestingly, 15 of the 25 identified proteins are LD-proteins from our database (with 9 of these confirmed in other LD databases). Several of these proteins have been previously implicated in AD pathogenesis. For example, the inflammatory transcription factor STAT3 and sorting nexin (SNX4) are of particular interest based on their contribution to inflammation and amyloid beta accumulation, respectively.^54-55^ Further investigation of how these proteins – and others identified here – contribute to LD-related function and AD pathology is warranted.

Another noteworthy overlap in the LD-proteome is with data from a large-scale proteomics analysis of AD postmortem tissue.^42^ There, Johnson et al. identified a network module (“M4”) of putative microglial metabolism proteins that shared ontology with microglial genes from an AD mouse model.^42^ Of these thirty M4 network proteins, we found 27 to be LD-enriched, and these proteins include previously established DAM markers such as Lgals3 and Fabp5^18^. While that particular study did not include *APOE* genotype in its analysis, a previous study from the Seyfried laboratory highlighted differentially expressed proteins from E4-expressing brain tissue.^43^ Again, our LD-enriched proteome showed high overlap, with around one third of the proteins up-regulated in the E4 AD (or E3 AD) brain also being upregulated on the surface of E4 (or E3) LDs. Finally, differentially expressed genes in microglia from E3- or E4-expressing mice treated with either saline or LPS^19^ also showed substantial overlap with our LD proteome, again highlighting the potential role of *APOE* and LDs within this critical immune cell. Interestingly, in each case above, gene ontology of these shared LD-AD proteins pointed toward a depletion of FA β-oxidation on E4 relative to E3 LDs. As from several previous studies, this highlights a potential pathogenic role of E4 in modulating glial FA metabolism^24-29^ with the data here implicating LDs as the initiating hub.

Though we performed lipidomics and proteomics analyses on LDs isolated from the liver rather than brain by necessity, this strong overlap with previous AD-microglia proteomic and transcriptomic datasets supported a translation of these findings to LD dynamics in this cell type. Related to this, a handful of studies have inhibited LD formation and observed rescue of some abnormal cellular activity as a result. For example, Marshallinger et al observed increases in pro-inflammatory cytokines, ROS, and abnormal function in LD-accumulating microglia (LDAM).^31^ When LDs were inhibited with Triacin C, the authors noted normalization in ROS, but did not comment on the restoration of cytokine output after LD inhibition. A similar observation has been made in E4 induced microglia-like (iMG) cells whose conditioned media suppressed the activity of neuronal cultures, whereas Triacin C sufficiently reduced LD load and mitigated this suppression.^27^ Another recent study showed that E4/4 iMGs had significantly greater LD accumulation compared with isogenic E3/3 iMGs, and restoration of cytokines was achieved with by lowering LD load through PIK3CA inhibition.^28^ However, this was thought to occur through increased levels of autophagic flux and not through inhibition of droplet production.

It was our original hypothesis that the exaggerated accumulation of LDs in E4 microglia drives increased cytokine release, and thus prevention of LD formation in these cells may alleviate this pro-inflammatory phenotype. However, although IFN-γ secretion appeared to be dampened by LD inhibition, ACAT and/or DGAT inhibitors otherwise had minimal or no effect on cytokine output. There could be several possible reasons for these results. First, with no ability to form LDs in which to sequester fatty acids, the cells are vulnerable to fatty acid toxicity which could itself cause an increase in cytokine release.^56^ Alternative approaches that focus on promoting LD turnover, rather than directly inhibiting their formation, could address this question and potentially provide a better approach. Second, it is possible that the increased LD accumulation and greater cytokine release in E4 microglia are simply independent phenomena – they occur in parallel, but do not share a mechanistic pathway.

Despite being largely unaffected by LD inhibition, we did observe an increase in cytokine output from E4 cells relative to E3 at baseline. However, when stimulated with LPS, E3 microglia had a more robust cytokine response, a finding that adds to a complicated literature. For example, Machlovi et al. previously showed that primary microglia expressing human E4 exhibit changes in morphology and increased cytokine production associated with LD accumulation compared to E3 microglia.^26^ Additionally, several *in vivo* studies of E4 mice and humans show an exaggerated response to inflammation in E4 individuals.^57-60^ Conversely, Kloske et al showed an overall reduction in inflammatory gene expression in human AD brains from E4 carriers compared to non-carriers.^61^ In vitro, primary astrocytes from E4 targeted replacement mice also had a reduced cytokine response after LPS stimulation compared to E3 astrocytes,^62^ while another study noted significant sex-based cytokine secretion differences in microglia from E3 and E4 targeted replacement mice.^63^ Thus, the impact of *APOE* variants across various cell types, the role of LDs in this process, and their integration to facilitate a coordinated immune response, all remain important areas of uncertainty for the field.

Our study has several limitations. First, due to the nature of the LD enrichment, some proteins that transiently associate with LDs may have been included in the analysis. This means that proteins that are not truly LD “resident” (i.e. wholly embedded within the LD phospholipid shell) are likely included in our analyses. However, because LDs transiently interact with many other organelles,^58^ we believe inclusion of these interacting partners provides important insight into potential *APOE*-dependent changes in LD dynamics within the cell. In fact, if/how *APOE* may mediate organelle-organelle interactions within the cell – for example LD-mitochondria or LD-lysosomal contacts – may dictate broader functional consequences and should be a focus for future study. Second, it should be noted that these results differ from recent experiments using similar methods in human induce pluripotent stem cells (hiPSC),^64^ where inhibiting TG biosynthesis in E4 microglia blunted cytokine/chemokine release and attenuated the disease-associated transcriptional profile of these cells. Differences in ApoE and human/mouse receptor binding affinity and interactions – and their downstream effects on lipid processing^65-67^ – may explain these varying results. Finally, while the current study leveraged the abundant and easily accessible LDs from the liver of human E3 and E4 expressing mice, it will be important to overcome the logistical hurdles of enriching LDs from the brain and microglia in order to conduct similar profiling of human cells and tissue.

In sum, our findings demonstrate that LD composition and dynamics are modulated by *APOE* genotype. Additionally, integration of our novel APOE-LD proteome with previous AD and microglia datasets, excitingly suggests that many of these previously implicated proteins may be LD-resident. Combined with a strong body of literature that supports a role for *APOE* in modulating metabolism, neuroinflammation, and LD formation, our work gives credence to the hypothesis that LDs may act as a primary hub in integrating the immunometabolic response of glia in AD. However, lipid droplets are essential for protection against lipotoxicity and for buffering ROS in the cell, thus preventing and inhibiting their formation may not be an optimal strategy. Instead, a more effective approach might involve modulating their regulation, such as by influencing their interactions, functions, or turnover. Therefore, additional studies focused on targeting LD functionality, or on shifting the protein profile of droplets toward a more neutral E3-like state – for example reprogramming E4’s metabolic state to a more anti-inflammatory aerobic production of ATP through oxidative phosphorylation – may hold promise for future AD therapy.

## METHODS

### Mouse Model

Targeted replacement mice expressing human *APOE3* and *APOE4* under the endogenous *APOE* mouse promoter were used in the following experiments.^68-69^ All mice received food and water ad libitum and were housed under a standard light/dark cycle. Procedures concerning mice were in compliance with the University of Kentucky’s Institutional Animal Care and Use Committee.

### Lipid Droplet Enrichment

Twelve-month-old female mice were used to determine LD dynamics at baseline and upon inflammatory stimulation between E3 and E4 genotypes. Mice were injected with saline or lipopolysaccharide (LPS) at 5mg/kg (n = 5 per group). After 24 hours, mice were humanely euthanized with a lethal injection of pentobarbital. Mice were perfused with saline and 300-400g of liver tissue was removed for lipid droplet enrichment. On ice, liver tissue was minced and bathed in homogenization buffer from Cell Biolabs Inc. (product No MET-5011). Homogenate was collected into a 2mL vial and 600ul of density gradient separation buffer from the LD kit was gently layered on top. Samples were spun at 13,500 rpm for 3 hours. A layer of lipid known as a fat-pad appeared at the top of the vial after centrifugation and this buoyant layer was enriched with lipid droplets. The top 270μl was removed, along with the fat pad, and placed into a new tube. Two liver sections per mouse were processed this way and flash frozen in liquid nitrogen to send for proteomics and lipidomics. An additional section of liver was centrifuged and the top 270μl was added to 1mL of ice-cold acetone for protein precipitation and western blot analysis.

### Proteomics Analyses

Proteomics: After lipid droplet enrichment via density gradient centrifugation, the top 270ul and ‘fat pad’ containing lipid droplets was removed and placed into an Eppendorf tube. Tubes were frozen in liquid nitrogen and placed in a -80C freezer prior to shipment to BGI Global for quantitative TMT proteomics analysis using methods and normalization procedures previously described.^68^ Briefly, two normalization procedures were applied for a 30-plex experiments (3 TMT experiments with 10 channels each). Within each 10-channel TMT run, the grand total reporter ion intensity for each channel was multiplied by scaling factors globally to adjust the channel intensity to the average total intensity across all ten channels in order to correct for sample loading and reaction efficiency differences. Second, common, pooled internal standards were utilized to normalize reporter ion intensities between the three TMT experiments.

Proteomics analysis detected 6,192 proteins. Proteins were accepted if they contained more than two unique peptides and were within the false discovery rate of 5%. A final 4,267 proteins were included and analyzed. These final proteins are considered as both LD-resident proteins and LD-interacting proteins. To determine which proteins may be true LD resident proteins, the accepted 4,267 were referenced against 6 known lipid droplet proteomes from liver cells and mouse and rat liver tissue.^21,36-39^ Upon cross-referencing our database with the six others, 2099 overlapping proteins were identified and considered as LD-resident proteins. Abundance averages were calculated within groups and multiple group t-tests were performed on the entire dataset. Significantly differentiated proteins were plugged into the open-source gene ontology analysis site, Enrichr, to understand protein interactions within biological processes. Volcano plots were made for each condition in the entire data set and the overlapping data set. Our full database was also compared against proteomes from papers investigating differences in the brain, Alzheimer’s disease, and microglia.^18,40-42^ We analyzed overlapping proteins from our database with these papers and further investigated their contribution to our hypothesis.

### Lipidomics analyses

After LD enrichment via density gradient centrifugation, the top 270μl and fat pad containing LDs was removed and placed into an Eppendorf tube. Tubes were frozen in liquid nitrogen and placed in a -80°C freezer. Samples were packaged and sent to BGI global. Nontargeted lipidomics analysis was performed using LC-MS/MS. High resolution mass spectrometer Q Exactive (Thermo Fisher Scientific, USA) was used for data acquisition in positive-ion and negative-ion mode respectively to improve the lipid coverage. The data were processed by LipidSearch 4.1 and BGI’s statistical software package, metaX. Lipidomic analysis revealed 321 unique lipid metabolites. An outlier analysis was performed in Prism Graphpad and revealed 176 statistical outliers among the 20 samples and 321 metabolites (2.7% outliers from original data). Metabolite abundance averages were calculated within groups and multiple group t-tests were performed on the entire data set.

### Oil Red O staining and analysis

Liver sections from mice treated with LPS or saline were stained with Oil Red O and Hematoxylin to quantify LD accumulation. Dissected liver specimens were fixed in 4% PFA for 24 hours then sunk in 30% sucrose + 0.05% sodium azide before being fixed in O.C.T. Free-floating sections were cut at 10μM and washed 3x for 5 minutes with PBS prior to staining. Sections were placed in 100% propylene glycol for 5 minutes then incubated in pre-heated Oil Red O (Abcam #150678) for 6 minutes at 60°C. After staining, sections were placed in 85% propylene glycol for 1 minute, rinsed with 2 changes of distilled water, and counterstained with Hematoxylin. Sections were mounted with ProLong Gold upon air drying. Liver sections were imaged on the Zeiss Axio Scan Z1 and 3 areas from each sample were randomly selected for droplet analysis. Total lipid droplet area was assessed using an automated algorithm which removed white (blank) background from the images and separated the colors to highlight the red-stained droplets. Individual droplet area was evaluated using ImageJ based on circularity of the droplets.

ORO images were loaded in a BGR format using the OpenCV module in Python. A Gaussian filter with a kernel value of 5 was applied to the images in order to reduce the noise of the pixels. The now blue lipid droplets were thresholded by performing color detection of the range of red (corresponding to the other tissue), using the cv2.inRange function and defining the boundaries in the RGB color space. The lower limit of the color was set as [60, 20, 2], and the upper limit, as [126, 255, 255]. A binary mask was then imposed on top of the original image, producing a new one where the values of the pixels within the defined color range become 0 (black). Upon counting the non-black pixels, that number was then multiplied by 100 and divided by the total amount of pixels conforming the image. The segmented images were saved into a database.

For individual lipid droplet area calculation, the segmented images were imported into ImageJ and subsequently turned into 8 bit grayscale images using the command Image>Type>8 bit (black and white). A new mask was imposed on top of the image using the command Image>Adjust>Threshold, setting the lower limit as 0 and the upper limit as 20 to re-cover the black pixels in the image. Upon inverting the colors of the image using the command Edit>Invert, the droplets were quantified and measured using the command Analyze>Analyze particles; the pixel size was set to 0-100 and circularity was set to 0 (1 being a perfect circle). Configurations were set to display the resulting outlines as new images. Perimeter and area measurements were selected and found under the menu Analyze>Set measurements. Individual droplet count and area was used to create a LD size histogram. Total LD area per image was calculated based on an area under curve analysis of this generated histogram.

### WGCNA

Weighted gene co-expression network analysis (WGCNA) (v1.70-3) was used to identify gene modules and build unsigned co-expression networks, including both negative and positive correlations. Briefly, WGCNA constructs a gene co-expression matrix, uses hierarchical clustering in combination with the Pearson correlation coefficient to cluster genes into groups of closely co-expressed genes termed modules, and then uses singular value decomposition (SVD) values as module eigengenes (MEs) to determine the similarity between gene modules or calculate association each module with sample traits (ex. *APOE* genotype and treatment). The top 3,000 variable genes were selected to identify gene modules and network construction. Soft power of 6 was chosen by the WGCNA function pickSoftThreshold. Next the function TOMsimilarityFromExpr was used to calculate the TOM (Topological Overlap Matrix) similarity matrix via setting power = 6, networkType = “signed”. The distance matrix was generated by subtracting the values from the similarity adjacency matrix by one. The function flashClust (v.1.01) was used to cluster genes based on the distance matrix, and the function cutreeDynamic was utilized to identify gene modules by setting deepSplit =3. Cytoscape (v.3.8.2) was applied for the gene-gene network visualization.

### Western Blotting

Protein from the lipid droplet enriched fraction was precipitated with acetone and resuspended in RIPA with protease inhibitor. The sample was diluted with 10 μl MilliQ water, and then further diluted at a 1:1 ratio with 2x Laemmli Sample Buffer (Bio-Rad Laboratories, Hercules, CA, USA). The samples were heated at 96.5° Celsius for 10 minutes, after which they were chilled on ice for 5 minutes before loading into the gel. 20 μl of the protein sample was loaded on 4-20% Criterion TGX Gels (Bio-Rad Laboratories, Hercules, CA, USA). Gels were transferred onto 0.2 μm nitrocellulose membrane (Bio-Rad Laboratories, Hercules, CA, USA) using the Trans-Blot Turbo Transfer System (Bio-Rad Laboratories, Hercules, CA, USA). After transfer, membranes were blocked for 30 minutes in 1% casein solution while gently rocking back and forth using the VWR Analog Rocker (Avantor, Radnor, PA, USA). The membranes were incubated overnight at 4° Celsius in 1:1000 PLIN-2 primary antibody solution (Novus Biologicals, Centennial, CO, USA). After incubation, the membranes were washed with PBS-T (0.05% Tween-20), three times for five minutes each wash. The membranes were then incubated for one hour at room temperature, while protected from light, in 1:5000 Goat α-rabbit IR 700 secondary antibody solution. (Bio-Rad Laboratories, Hercules, CA, USA). They were then washed with PBS-T, three times for five minutes each, and then with PBS two times for five minutes each. Membranes were imaged using a ChemiDoc XRS Imaging System (Bio-Rad Laboratories, Hercules, CA, USA).

### Cell Culture

Human *APOE* expressing mice were bred and pups were taken at postnatal day P1-3 to obtain primary microglia for experiments. Littermates were pooled together to increase cell counts, but pups from the same litter were not genotyped for sex chromosomes. During dissection, forebrains were removed from mice and placed in a dissection buffer consisting of ice-cold Hanks Balanced Salt Solution (HBSS #). Under the dissection scope, meninges were carefully removed and the brain was placed in DMEM F12 media (10% FBS and 1% pen/strep). Pooled brains were cut into small pieces and digested with trypsin at 37C for 25 minutes, agitating every five minutes. Trypsin was neutralized with an equal volume of DMEM media and brains were centrifuged for 5 minutes at 400xg prior to washing with HBSS. Brain homogenate was resuspended in DMEM, triturated, and ran through a 70 micron filter. Mixed glial cells from this preparation were split into T75 flasks (1 flask per brain). Dead cells and debris were washed after day 1 in culture and media was switched to a supplemented DMEM: F12 media on day 7. After two weeks, microglia begin to emerge from the glial cells. Flasks were placed on an orbital shaker for two hours and media was collected and centrifuged to pellet the microglia. Cells were resuspended in DMEM, counted, and plated for various experiments.

### Lipid Droplet Imaging

For lipid droplet imaging experiments, cells were seeded at 60,000 cells per well on a poly-L-lysine coated 12mm glass coverslip. Cells were treated with 250uM oleic acid conjugated with BSA, 10ng lipopolysaccharide (LPS), necroptotic Neuro-2A cells (nN2A), N2A plus LPS, oleic acid plus LPS, or an untreated control. After 24 hours, media was collected for cytokine analysis and cells were fixed in 4% paraformaldehyde. Cells were stained with BODIPY and coverslips were mounted with a DAPI nuclear staining mounting media. N2A cells were split into a few dozen T-75 flasks with non-filter cap lids. Once cells grew to confluency, the caps were tightened to induce hypoxia and cell death. Cells began to detach from the flasks and all apoptotic cells and media were collected, split into 8×10^6^ cells/mL aliquots and frozen for later use. Microglia were treated at a 1:10 ratio of microglia to nN2A cells during experiments.

Imaging took place on a Nikon A1R confocal microscope. 405nm wavelength was used to acquire microglial nuclei stained with DAPI and 488nm wavelength was used to acquire neutral lipid stained with BODIPY. Color threshold analysis using ImageJ was used to select high intensity regions of neutral lipid, indicating the presence of lipid droplets. Lipid droplets were then quantified by measuring the threshold area.

### Cytokine Analysis

Primary microglia from E3 and E4 targeted replacement mice were plated in 96-well plates and allowed to adhere. For a 24-hour treatment, cells were treated with either a DMSO control, DGAT inhibitor (10μM A9250000), ACAT inhibitor (10μM Avisimibe), or both inhibitors. Six conditions were tested: control, 250μM OA, 10ng LPS, 1:10 ratio of nN2A cells, OA + LPS, and nN2A + LPS. Cytokines were measured using a custom V-plex assay from MesoScale Discoveries with probes for IL-1ß, INGy, TNF, IL-10, and IL-6.

## Author Contributions

CMF, JMM, SMG, SC and LAJ designed the experiments. CMF, IOS and LAJ analyzed the data, generated figures, and wrote the paper. LAJ and JMM supervised the ‘omics analysis with data analysis by CS, SL and GMS. NAD assisted with primary cell culture and SMG assisted with lipid droplet isolation with technical assistance from CMF. All authors read the paper and provided edits.

## Acknowledgements

We would also like to thank Darcy Adreon, Gabriella Morillo-Segovia, and Drs. Jessica Funnell and Gabriela Hernandez for their invaluable assistance.

