## Supplementary Figure for "*APOE4* alters the lipid droplet proteome and modulates droplet dynamics"

### Supplemental Figures

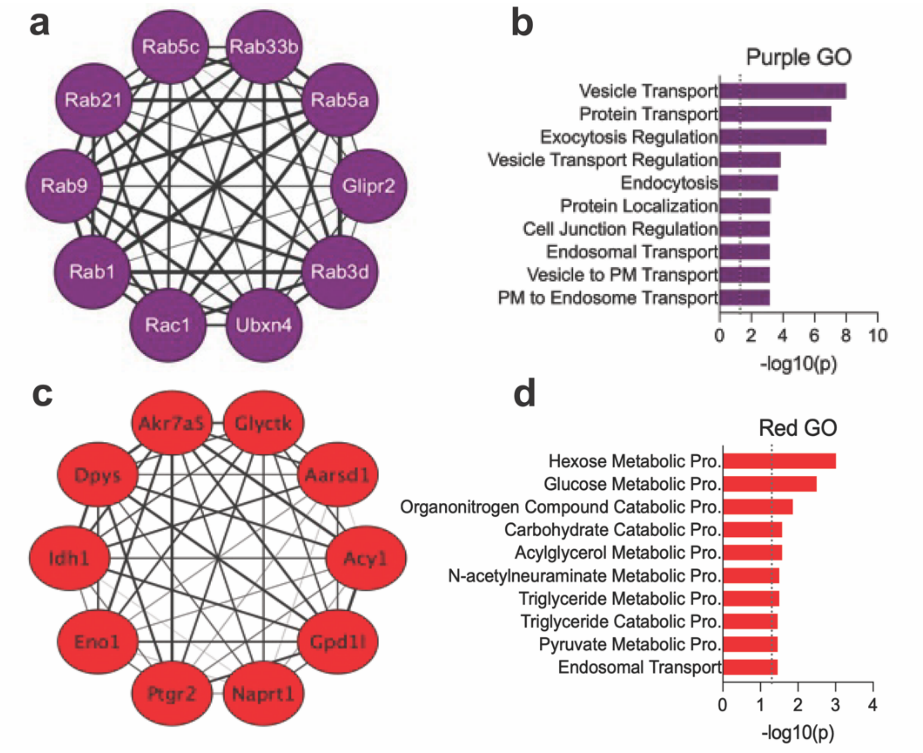


**Supplement Figure 1.** (**a**) The purple module was enriched in proteins involved in vesicle transport and was significantly different in the LPS groups compared to saline treated in both E3 and E4. The 10 ten proteins from this module are displayed in the circle plot along with gene ontology form the entire module. (**b**) The red module was enriched in pathways involved in carbohydrate metabolism and was lower in both LPS groups when compared to saline treated. The top 10 genes from this module are displayed in the circle plot along with the gene ontology form the entire module.

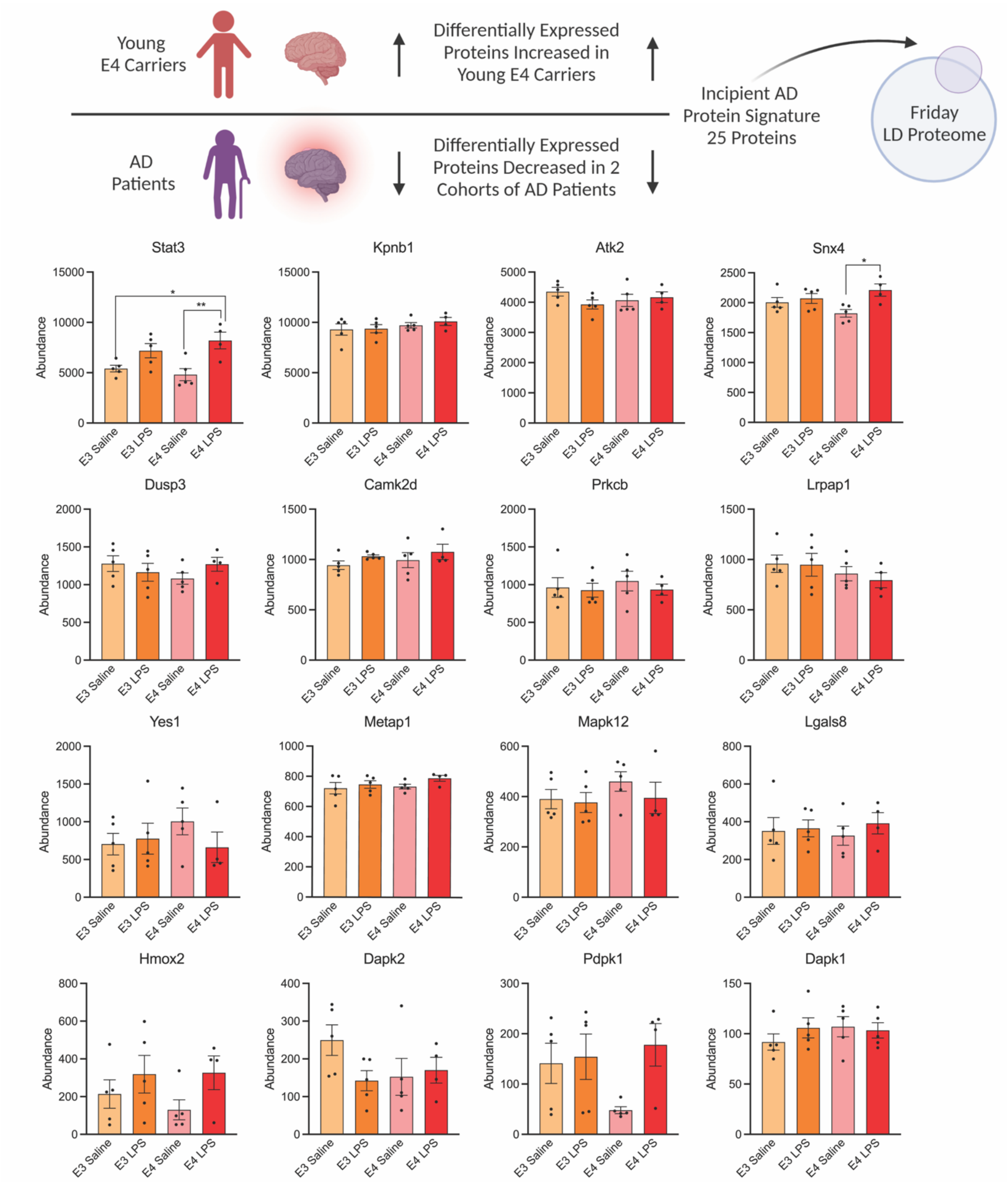


**Supplement Figure 2.** This “incipient AD” (iAD) signature showed 60% overlap with the APOE LD proteome from the current study. Shown are the individual graphs for each of the overlapping proteins. *p<0.05, **p<0.005, two-way ANOVA analyses

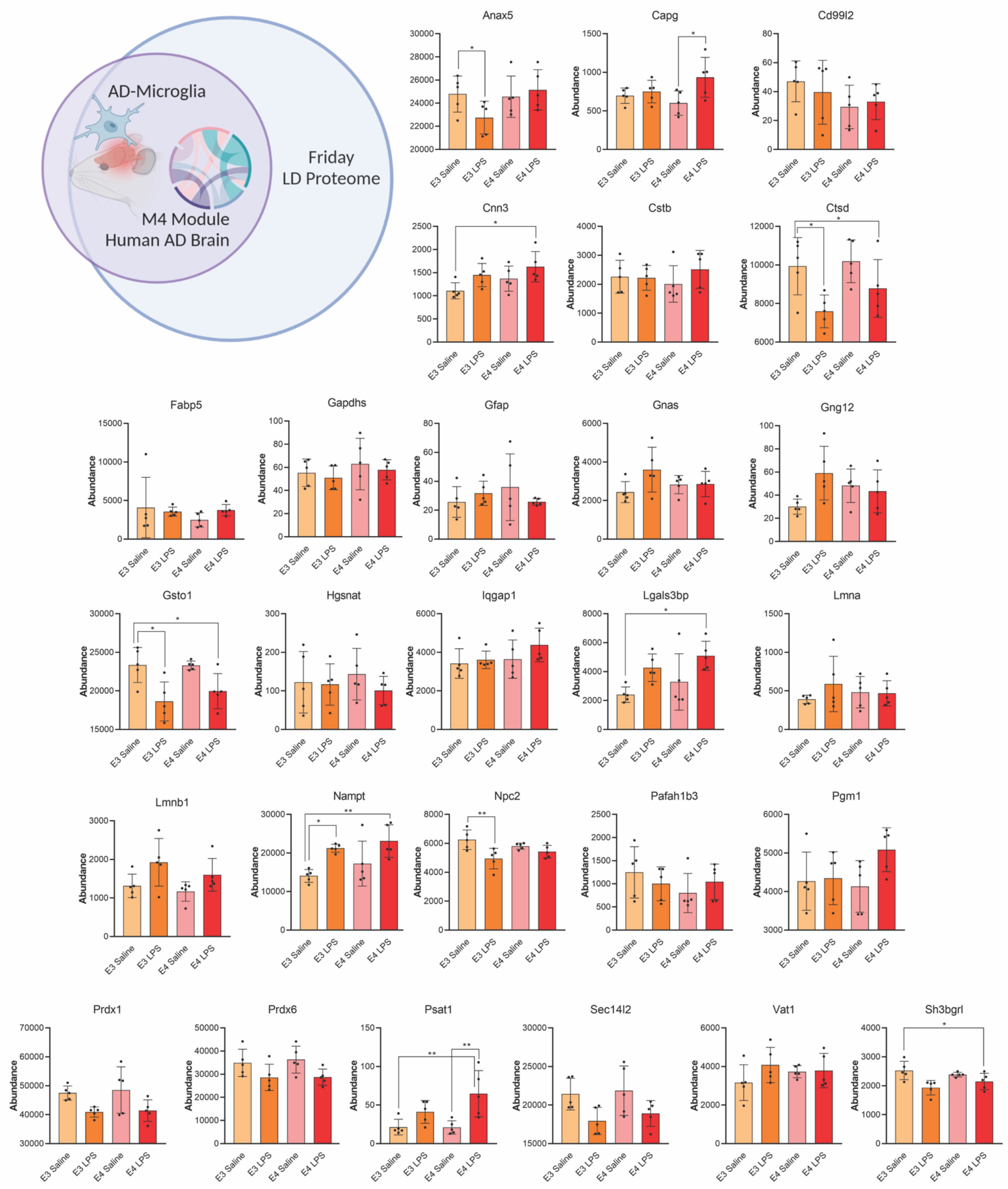


**Supplement Figure 3.** A large proteomic study by Johnson et al. revealed a glial metabolism module ("M4") which was differentially expressed in AD brains as well as in microglia from a mouse model of AD. This AD-glial-metabolism signature has 90% overlap with our LD proteome. Shown are the individual graphs for each of the overlapping proteins. *p<0.05, **p<0.005, two-way ANOVA analyses.

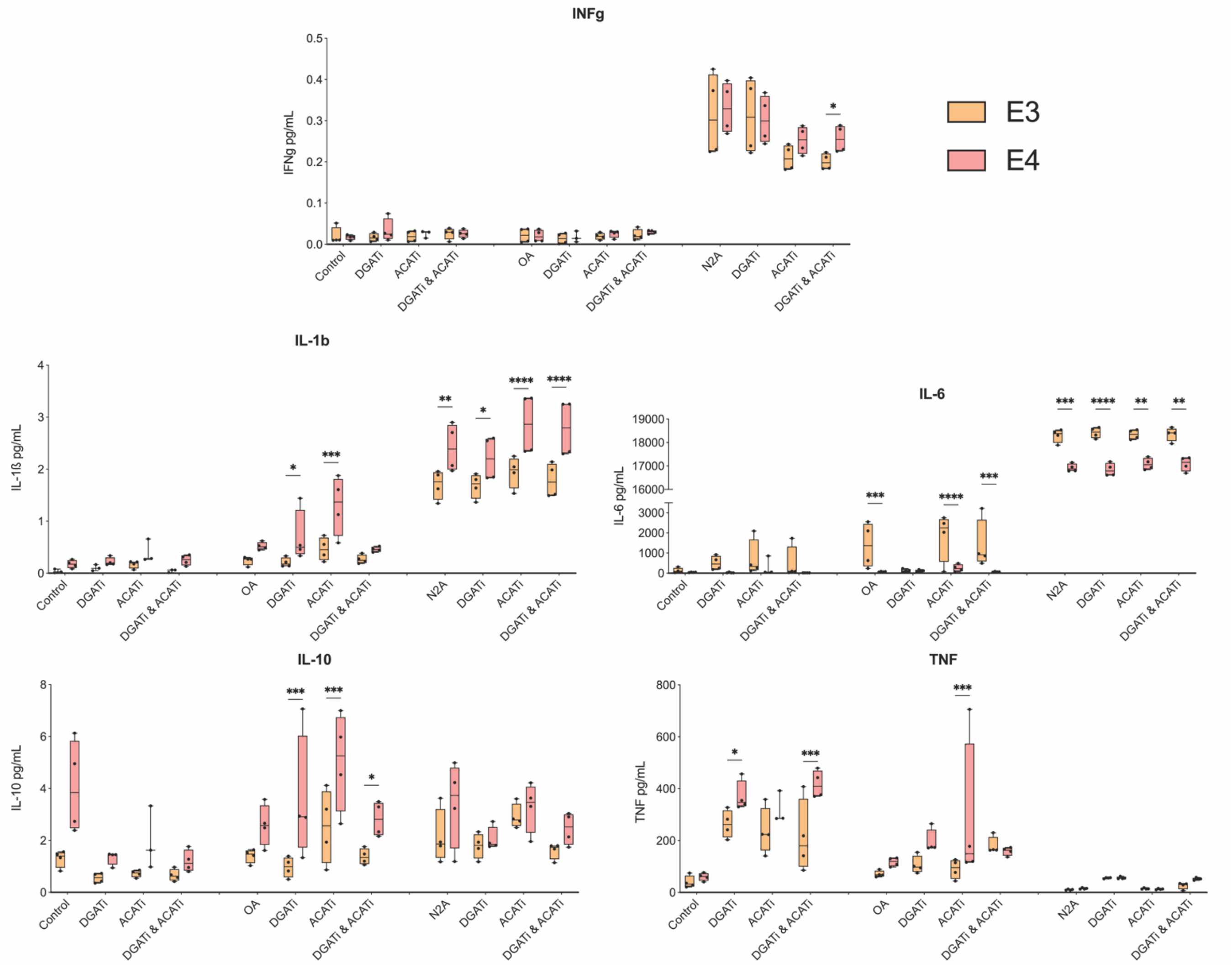


**Supplement Figure 4.** Concentrations of INFg, IL-1ß,IL-6 TNF, IL-10, and TNF measured in the media of E3 or E4 microglia. Microglia were given either vehicle, DGATi, ACATi, or both plus loading with OA or nN2A. *p<0.05, **p<0.01, ***p<0.001, ****p<0.0001 two-way ANOVA analyses.

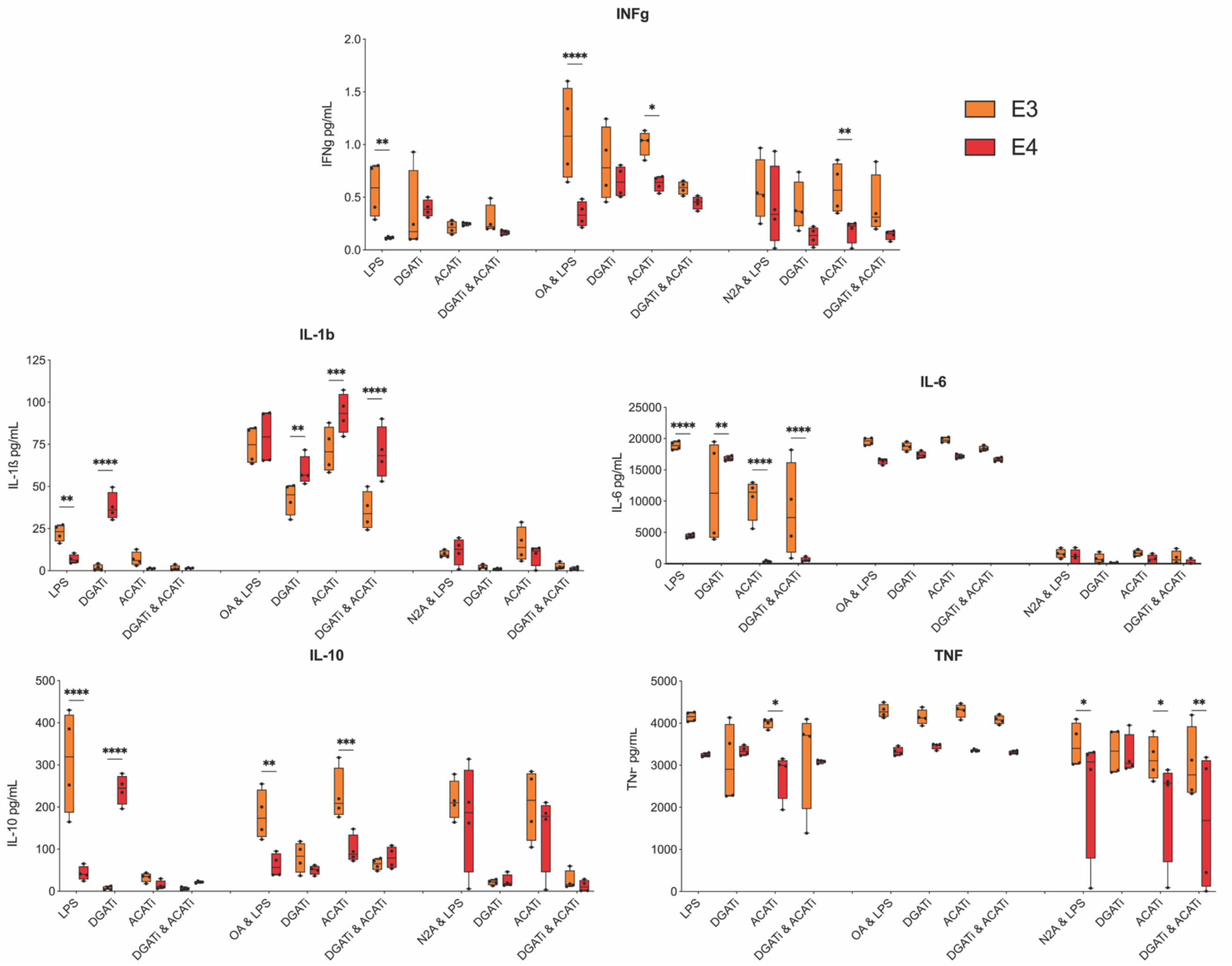


**Supplement Figure 5.** Concentrations of INFg, IL-1ß,IL-6 TNF, IL-10, and TNF measured in the media of LPS treated E3 or E4 microglia. Microglia were given either vehicle, DGATi, ACATi, or both plus loading with OA or nN2A. *p<0.05, **p<0.005, ***p<0.001, ****p<0.0001 two-way ANOVA analyses.
